# Amplified visual immunosensor integrated with nanozyme for ultrasensitive detection of avian influenza virus

**DOI:** 10.1101/128538

**Authors:** Syed Rahin Ahmed, Suresh Neethirajan

**Affiliations:** BioNano Laboratory, School of Engineering, University of Guelph, Guelph, Ontario, Canada N1G 2W1

**Keywords:** Dual color enhancement, Avian influenza virus detection, Gold nanoparticles, Peroxidase mimic

## Abstract

Nanomaterial-based artificial enzymes or nanozymes exhibits superior properties such as stability, cost effectiveness and ease of preparation in comparison to conventional enzymes. However, the lower catalytic activity of nanozymes limits their sensitivity and thereby practical applications in the bioanalytical field. To overcome this drawback, herein we propose a very simple but highly sensitive, specific and low-cost dual enhanced colorimetric immunoassay for the detection of avian influenza virus A (H5N1) through facile *in situ* synthesis of gold nanoparticles and their peroxidase-like enzymatic activity. 3,3’,5,5’-Tetramethylbenzidine (TMBZ) was used as a reducing agent to produce gold nanoparticles (Au NPs) from a viral target-specific antibody-gold ion complex. The developed blue color from the sensing design was further amplified through catalytic activity of Au NPs in presence of TMBZ–hydrogen peroxide (H_2_O_2_) complex. The developed dual enhanced colorimetric immunosensor enables the detection of avian influenza virus A (H5N1) with a limit of detection (LOD) of 1.11 pg/mL. Our results further confirms that the developed assay has superior sensitivity than the conventional ELISA method, plasmonic-based bioassay and commercial flu diagnostic kits.

## 1. Introduction

Colorimetric detection of infectious virus pathogens based on specific antigen and antibody interaction enables the development of a low-cost, field-portable sensor, and has a tremendous ability to identify target analytes at an early stage in complex biological matrices (Xu *et al*., 2017; Rakow *et al*., 2000; Song *et al*., 2011; Ye *et al*., 2015). Compared to the conventional detection techniques such as electrochemical, fluorescence or surface plasmon resonance; colorimetric visualization of disease-related biomarker analytes has drawn much attention recently due to its simplicity and practicality (Gao *et al*., 2014a).

To generate significant naked-eye detectable color, natural enzymes are widely used as a labeling agent with antibodies. For example, horseradish peroxidase (HRP) and alkaline phosphatase (ALP) labeled immunoreagents are commonly used in the colorimetric immunoassay (Gao *et al*., 2104a; Malashikhina *et al*., 2013). However, denaturation, low stability (temperature and pH), high cost and complex purification methods hamper their use in practical field applications.

Recently, advances in nanotechnology have permitted the alternation of natural enzymes and overcome their drawbacks, i.e., the use of nanomaterials instead of natural enzymes. So far, nanomaterial-based artificial enzymes have rapidly been emerging as a research interest in the nanobiotechnology field, and their applications have been extensively explored in bioanalysis, bioimaging and biomedicine (Weiz *et al*., 2013). Nanomaterials such as Fe3O4 magnetic nanoparticles (MNPs), platinum nanoparticles (Pt NPs), cerium oxide nanoparticles (CeO2 NPs), gold nanoparticles (Au NPs), copper oxide nanoparticles (CuO NPs), carbon nanotubes and graphene possess enzyme-like activities (Gao *et al*., 2007b; Polsky *et al*, 2006; Asati *et al*., 2009; Ahmed *et al*., 2016a; Chen *et al*., 2012; Ahmed *et al*., 2016b; Ahmed *et al*., 2017c). Au NPs have frequently been used for the construction of diverse colorimetric biosensors because of their intriguing size-dependant optical properties, ease of preparation, stability and strong catalytic activities. These properties make them a promising nanomaterial for various applications (Ahmed *et al*., 2016b; Ahmed *et al*., 2017c; Jv *et al.;* 2010).

However, the catalytic activity of nanozymes is lower than that of their natural enzymes, which ultimately effects sensitivity of the analyte detection and limits their use in bioanalysis (Cheng *et al*., 2016). To address these challenges, efforts have begun to develop nanozymes with enhanced catalytic activities. An enzyme-cascade-amplification strategy opens a new horizon in bioassay where natural enzymes and nanozymes work together within a confined environment.^5^ This cascade reaction is catalyzed by nanozymes and results in a signal amplification of several orders of magnitude within milliseconds.

Nanohybrids that contain two or more nanomaterials in a single entity has the potential to enhance the enzymatic activities of nanozymes. The poor catalytic activities of individual nanoparticles and low dispersibility of some nanomaterials (eg: CNTs, graphene) could partially be overcome by combining various nanomaterials into one superstructure (Ahmed *et al*., 2017c). Such nanohybrids possesses higher enzymatic activity and are suitable to develop ultra-sensitive colorimetric immunosensor.

Despite the significant progress in terms of the catalytic activity of nanozymes, cost-effectiveness due to the use of different chemicals and multistep preparation processing remains a weakness. This problem continues to impede the use of nanozymes in real-life applications. In addition, the lack of monodispersibility and uniform size of nanomaterials might be affecting the catalytic activities. Therefore, new strategies are urgently needed to design an applicable colorimetric detection system considering the aforementioned limitations.

In this study, a dual enhanced colorimetric detection system for avian influenza A (H5N1) virus was designed using lower amounts of chemicals and by avoiding a multistep synthesis process. The proposed detection strategy used an *in situ* synthesis of Au NPs from a target virus–specific antibody and gold ion complex using TMBZ. At this stage, a blue color was developed due to oxidization of TMBZ, which deepened in color upon addition of a TMBZ-H_2_O_2_ system due to the catalytic activity of synthesized Au NPs. The developed synthesis process does not require washing steps or the modification of the enzymatic activities of Au NPs. The change of color is directly correlated with the virus concentration, and hence enables the monitoring of color changes in naked-eye to determine the presence of the target avian influenza virus in the sample.

## 2. Experimental Section

### 2.1. Materials and reagents

Gold (III) chloride trihydrate (HAuCl_4_·3H_2_O) 3,3’,5,5’-tetramethylbenzidine (TMB), Hydrogen peroxide (H_2_O_2_), Nunc-Immuno 96-well plates were purchased from Sigma-Aldrich (St. Louis, MO, USA). The anti-influenza A (H5N1) virus hemagglutinin (HA) antibody [2B7] (ab135382, lot: GR100708-16), recombinant influenza virus A (Avian/Vietnam/1203/04) (H5N1) (lot: GR301823-1), goat anti-mouse IgG, horseradish peroxidase (HRP)-conjugated whole antibody (Ab 97023, lot: GR 250300-11) and immunoassay blocking buffer (Ab 171534, lot: GR 223418-1) were purchased from Abcam, Inc. (Cambridge, UK). Recombinant influenza virus A (H1N1) (California) (CLIHA014-2; lot: 813PH1N1CA) was purchased from Cedarlane (Ontario, Canada). Influenza A (H5N2) hemagglutinin antibodies (Anti-H3N2 antibodies HA MAb, Lot: HB05AP2609), Influenza A (H7N9) hemagglutinin antibodies (Anti-H7N9 antibody HA MAb, Lot: HB05JA1903), recombinant influenza virus A (H5N2) HA1 (A/Ostrich/South Africa/A/109/2006)(lot: LC09AP1021), recombinant influenza virus A (H7N8) HA1 (A/Mallard/Netherlands/33/2006) (lot: LC09AP1323) and recombinant influenza virus A (H7N9) HA1 (A/Shanghai/1/2013) (lot: LC09JA2702) were purchased from Sino Biological, Inc. (Beijing, China). All experiments were performed using highly pure deionized (DI) water (>18 MΩ· cm).

### 2.2. Preparation of antibody and gold ion complex

The bioconjugates of anti-HA H5N1 antibodies (Ab 135382) with gold ion (Au^3+^) were prepared based on electrostatic interaction as follows: 1 μg/mL (1 mL) antibody solutions were prepared in phosphate-citrate buffer solution in which 5 mM (60 μL) HAuCl_4_ solution was added and gentle stir for 30 min at 4500 rpm speed (Southwest Science, NJ, USA). The, antibody-gold ion complex was separated through high speed centrifugation (15000 rpm for 15 min) using D3024 Micro-centrifuge (DEELAT, Calgary, Canada) and redispersed in phosphate-citrate buffer solution. A detailed pH-dependent study was performed to check the stability of antibodies. We had not observed any changes in the solution color (light yellow) at room temperature for several months, indicating that the metal oxidation state Au (III) was unchanged.

### 2.3. Characterization of antibody specificity

The specificity of the anti-H5N1 HA (Ab 135382) for influenza virus A/ Vietnam 1203/04/2009 (H3N2) was evaluated using the ELISA technique. Briefly, viral stocks were diluted with phosphate-buffered saline (PBS, pH 7.5) to a final concentration of 1μg/ml to perform ELISA. Virus solution (100 μl) was then added to each well of a nonsterile polystyrene 96-well flat-bottom microtiter plate for overnight at 4°C to allow adsorption of the virus onto the plates. The plates were then rinsed with PBS (pH 7.5), and the surface pores were blocked with 100 μl of immunoassay blocking buffer (Ab 171534) for 2 h at room temperature. After rinsing three times with PBS (pH 7.5) solution, anti-H5N1 HA Ab (1 μg/ml), anti-H5N2 HA antibody (1 μg/ml) and anti-H7N9 HA antibody (1 μg/ml) (100 μl/well) were added to each of the wells, and the plate was incubated for 1 h at room temperature. After rinsed three times with PBS (pH 7.5), HRP-labelled secondary antibody was added to each well for 1h at room temperature. Again rinsed with PBS buffer and TMBZ (10 nM)/H_2_O_2_ (5 nM) solution were added to each wells (50 μL/well). 10% H_2_SO_4_ solution were added to each well (50 μL/well) to stop enzymatic reactions and the absorbance of the enzymatic product at 450 nm was measured using a microplate reader (Cytation 5, BioTek Instruments Inc., Ontario, Canada) to quantify the interaction of the antibodies with the influenza viruses.

### 2.4. Colorimetric detection of avian influenza A (H5N1) virus

Avian influenza virus A (H5N1) stock solution (0.47 mg/mL) was serially diluted with a PBS buffer (pH 7.5) to create sensing subjects for this experiment. Virus solution (100 μL) was then added to each 96-well plate and incubated for overnight at 4°C. BSA (100 μL,1ng/ml) was used as a negative control for testing the selectivity and specificity of the proposed sensing method. A series of different influenza viruses were used for evaluation of the specificity in this experiment. After rinsing with PBS (pH7.5) three times and being blocked with 100 μL of 2% skim milk for 2 h at room temperature, 50 μL of antibody-gold ion bioconjugates was added to the pre-adsorbed wells, and the plates were incubated for 1h at room temperature. After washing three times with PBS (pH 7.5), a 50 μL mixture of TMBZ (10 nM) was added into each of the wells for the color development (bluish-green color), and intensity was further developed with the addition of 50 μL TMBZ (10 nM)/H_2_O_2_ (5 nM) solution within seconds. Finally, 50 μL of 10% H_2_SO_4_ was added to each well to stop the reaction. The absorbance at 450 nm was measured, and a dose-dependent curve was constructed based on the absorbance values at different concentrations of avian influenza virus.

### 2.5. A comparison study between the proposed sensing system and the commercial ELISA kit

A comparison study was performed with the commercial avian influenza A H5N1 ELISA kit (Cat. No: MBS9324259, MyBioSource, Inc., San Diego, USA) to validate the proposed method. Different virus titers were prepared using sample diluent provided in the ELISA kit box and by strictly following the manufacturer’s protocol in the performance of the bioassay. Positive and negative avian influenza diagnostic results were obtained from different intensity of colors that appeared on the 96-well plates at room temperature.

### 2.6. Spectroscopy and structural characterization

Transmission electron microscopy (TEM) images were acquired using Tecnai TEM (FEI Tecnai G2 F20, Ontario, Canada). The ultraviolet-visible (UV–vis) spectrum was recorded using a Cytation 5 spectrophotometer (BioTek Instruments, Inc., Ontario, Canada). Antibody-gold ion conjugates were monitored by Fourier transform infrared spectroscopy (FT-IR spectroscopy) (FT/IR6300, JASCO Corp., Tokyo, Japan). Zeta potential was measured with Zetasizer Nano ZS (Malvern Instruments Ltd., Worcestershire, UK).

## 3. Results & discussion

### 3.1. In situ synthesis of gold nanoparticles and their catalytic activity study

In this study, TMBZ was chosen as a key chemical to synthesize Au NPs with bluish-green colored solution, as schematically depicted in **Figure 1**. We chose TMBZ for several reasons. Firstly, it acts as a reducing agent of gold ions and a stabilizer of Au NPs, forestalling the need for an extra stabilizer. Secondly, TMBZ can produce Au NPs with a positive charge (because of an –NH_2_ group in its chemical structure). These positively charged NPs have strong catalytic activity and color-producing ability towards the TMBZ-H_2_O_2_ complex compared to negatively charged Au NPs, as we reported earlier (Ahmed *et al*., 2016a). Therefore, synthesized Au NPs with bluish-green colored solution were further showed peroxidase-like activity and produced deeper colored solution by adding the TMBZ-H_2_O_2_ complex without any washing steps.

**Fig. 1.**
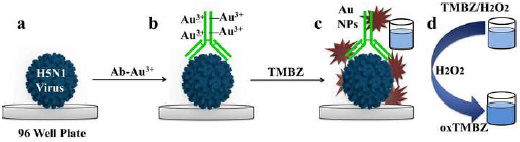
Schematic presentation of dual enhanced colorimetric detection. (a) Deposition of avian influenza virus on 96-well plates. (b) Target virus–specific antibody-gold ion added on virus surface. (c) Upon the addition of TMBZ, Au nanostructures forms, causing development of a bluish-green color. (d) The bluish-green color becomes more intense with the addition of a TMBZ–H_2_O_2_ solution due to the enzymatic reaction of the Au nanostructure.

Before starting the sensing assay, an experiment was performed with only gold ions to check the viability of our proposed method. The results are shown in Figure 2. It was clearly observed with the naked eye that a bluish-green colored solution was developed by the reaction of gold ions and TMBZ (Step 1, Fig. 2A-a), which in turn became more intense in color with the addition of the TMBZ–H_2_O_2_ complex (Step 2, Fig. 2A-b). Spectroscopic study of colored solutions (Step 2) revealed a several-fold enhanced spectrum peak centered at 655 nm compared to Step 1. In addition, a plasmonic peak with a broader spectrum of positively charged Au NPs (+25.6 eV) appeared at around 550 nm (Fig. 2B).

**Figure 2.**
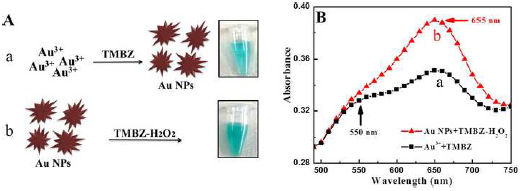
Viability test of proposed method. (A) The TMBZ reacted with the gold ions to form Au NPs with a bluish-green color. (a) The solution’s color deepened with the addition of the TMBZ-H_2_O_2_ solution. (b) (Inset: naked-eye image) (B) UV-visible spectrum of Steps (a) & (b).

### 3.2. Characterization of gold nanoparticles (Au NPs)

In the present work, the reducing ability of TMBZ was applied to prepare gold nanostructures by reducing gold chloroauric acid at room temperature. A charge-transfer complex (TMB and TMB^2^+) nanofibers due to the oxidation of TMB was also obtained along with gold nanostructures (Yang *et al*., 2008). A series of characterization was done to confirm the presence of a Au nanostructure. As shown in Figure 3, a TEM image revealed several micro-length TMB fibers with nanoscale diameter, which were formed during the reaction (Fig. 3A). A close view of the nanofibers shows nanoscale gold particles present inside the TMB fibers (Fig. 3B). The diameter of the nanostructured gold was approximately 50 nm with a non-spherical shape (Fig. 3C). The elementary analysis also confirmed the presence of large-scale TMB nanofiber and Au nanostructure (white dotted) in solution. In figure 3D, the elementary analysis confirmed the presence of a gold nanostructure with 72.4 and 31.73 weight and atomic percentage respectively. The crystal structure of the Au nanostructure was characterized using X-Ray Diffraction (XRD) analysis. The diffraction peaks appeared at 2θ values of 38.3, 44.1, 64.1, 79.2 and 81.6, representing (111), (200), (220), (311) and (222) planes of Au nanostructure respectively (Fig. 4A) (Ahmed *et al*., 2017c).

**Figure 3.**
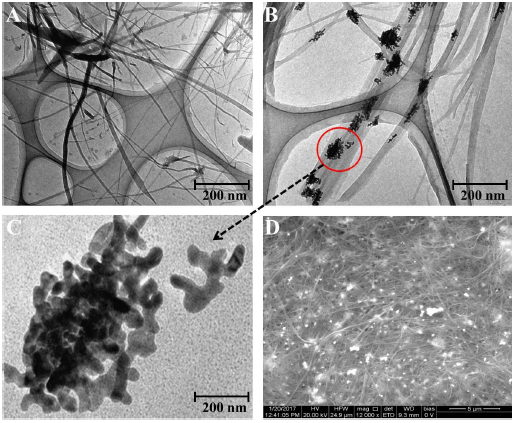
Microscopic study of TMBZ fiber and Au NPs. (A) TEM image of TMBZ fiber. (B) Closed view of TMBZ fiber. (C) Au nanostructure. (D) SEM image of TMBZ fiber and Au NPs.

### 3.3. Specificity of antibody towards target virus and its binding confirmation with gold ions

The specificity of anti-influenza A (H5N1) virus hemagglutinin (HA) antibody Ab 135382 against recombinant influenza virus A (Avian/Vietnam/1203/04) (H5N1) was confirmed using a conventional ELISA method. Fig. 4B shows that the optical density obtained for the target virus/Ab 135382/HRP-conjugated secondary antibody/ TMBZ–H_2_O_2_ complex was higher than those of Anti-H5N2 HA and anti-H7N9 HA Ab, implying the specificity of Ab 135382 against recombinant influenza virus A (Avian/Vietnam/1203/04) (H5N1).

**Figure 4.**
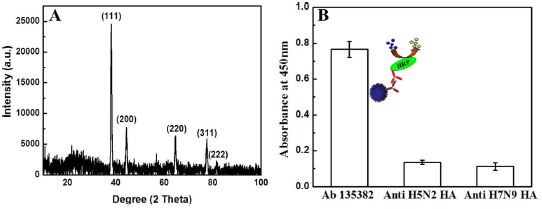
Optical study of Au nanostructure and antibody specificity towards target virus. (A) X-ray diffraction spectra of Au nanostructure. (B) ELISA results of antibody specificity towards target virus.

### 3.4. Binding confirmation of anti-H5N1 Ab 135382 with gold ions

The stability of anti-H5N1 Ab 135382 on HAuCl_4_ solution and its binding results with gold ions is shown in Figure 5. Here, Au^3+^ ions bind with negatively charged antibodies (-1.76 eV) through electrostatic force. At pH > 2, antibodies became aggregated and may lose their bio function. To avoid aggregation, a pH value ∼4 was kept during conjugation (Fig. S1A). The binding of antibodies and gold ions (Ab-Au^3+^) complex was confirmed by FTIR analysis. In Figure S1B, two new peaks arose compared to bare antibodies and HAuCl_4_ in the range of 3200-3000 cm^−1^, indicating new bonding between antibodies and HAuCl_4_. Also, higher optical density obtained for Ab-Au^3+^ complex in ELISA results further confirmed the successful binding between antibodies (Ab) and gold ions (Au^3+^) (Fig. S2).

**Figure 5.**
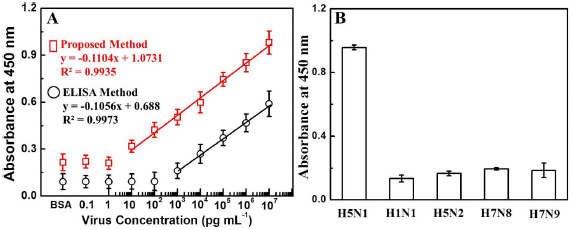
Detection of avian influenza virus A (H5N1). (A) The calibration curve of the absorbance corresponding to the concentration of avian influenza virus A (H5N1). BSA was used as a negative control. Squares (red line) and circles (black line) denote proposed and conventional ELISA sensing results, respectively. (B) ELISA results for selectivity of the present study with different influenza viruses. Error bars in (A) and (B) denote standard deviations (n=3).

### 3.5. Specificity of proposed bioassay

The catalytic activity of the proposed bioassay was investigated using four different reaction mixtures: a) recombinant influenza virus A (Avian/Vietnam/1203/04) (H5N1) /Ab135382–gold ion conjugates/TMBZ; b) replacing the specific Ab135382–gold ion conjugates with another anti-H5N2 HA–gold ion conjugate in the reaction mixture; c) removing the TMBZ from the reaction mixture and d) removing the gold ions (only Ab 135382) from reaction the mixture (a). A deep bluish-green color developed in the sample mixture (only (a)), and a strong characteristic absorption peak at 655 nm was also observed (Fig. S3). However, no such characteristic peak was observed for other mixtures (b, c and d), suggesting that the proposed sensing method is highly specific and color development occurs only with the target virus, its specific Ab-conjugated gold ions and TMBZ.

### 3.6. Quantitative analysis of avian influenza virus

A wide range of quantitative analysis for recombinant influenza virus A (H5N1) detection was performed after confirming the specificity and binding of Ab 135382 towards the target virus. The sensitivity of this proposed system for recombinant influenza virus A (H5N1) detection was found to be in the range from 10 pg/mL to 10 μg/mL with an LOD value of 1.12 pg/mL. In case of conventional ELISA method, the LOD value was calculated to be 909 pg/mL, suggesting that the proposed method was 811 times more sensitive than the conventional ELISA method (Fig. 5A & S4).

It is instructive to do a comparative study with other nanotechnology-based analytical techniques. To do that, we have compared the present technique with the plasmonic resonance peak response of synthesized Au NPs (at the intermediate stage) with different concentrations of target analytes. As shown in Figure S5, the change of plasmonic peak located at 550 nm was not consistent compared to the peak at 655 nm (present study) and showed 10 times less sensitivity (100 pg/mL) than the current detection technique.

To evaluate the selectivity for the detection of avian influenza virus, the proposed bioassay was implemented with other virus strains, namely H1N1, H5N2, H7N8 and H7N9. As shown in Fig. 5B, a significant change in the absorbance density (8~9-fold higher) was observed with the target avian influenza virus A (H5N1) in comparison to other viruses, revealing that the developed dual enhanced immunoassay was sufficiently selective for the detection of target avian influenza virus A (H5N1).

### 3.7. Comparison study of detection with commercial kit

The sensitivity of the proposed method was compared with those of a commercially available avian influenza A (H5N1) diagnostic kit (Table 1, Fig. S6). The naked-eye color response to the detection of avian influenza A (H5N1) in the commercial kit was as high as 1 ng/mL, indicating that our system is more sensitive than the commercial kit. Thus, the dual enhanced colorimetric technique presented here for visual detection of avian influenza viruses could be applicable for low-cost, visible, point-of-care diagnosis and also extendable to develop other nanozyme-based biomarker detection.

**Table 1:**
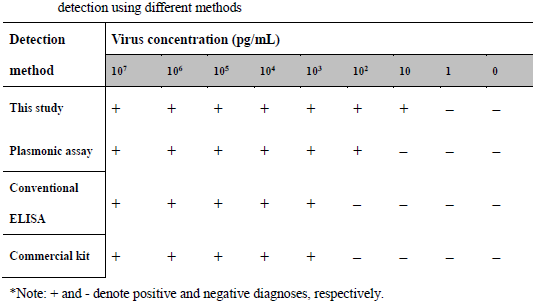
Comparison of avian influenza virus A (Avian/Vietnam/1203/04) (H5N1)

## 4. Conclusions

A facile design of enhanced colorimetric detection for avian influenza virus was developed in this study. A remarkable improvement of the color visualization, as well as detection sensitivity, was observed due to the dual effects of the sensing mechanism. The blue colour developed during the Au NPs synthesis process was further amplified bedue to the peroxidase-like activity of Au NPs in the presence of a TMBZ–H_2_O_2_ mixture. Our study not only provides a facile synthetic route for Au NPs but also opens an innovative approach for biosensor development using Au NPs as nanozymes.

## Acknowledgements

The authors sincerely thank the Natural Sciences and Engineering Research Council of Canada (400929), Ontario Ministry of Agriculture, Food and Rural Affairs (298634), Canadian Poultry Research Council (300142) and the Egg Farmers of Canada (201928) for funding this study.

## Appendix A. Supplementary material

Supplementary data associated with this article can be found in the online version at:

---

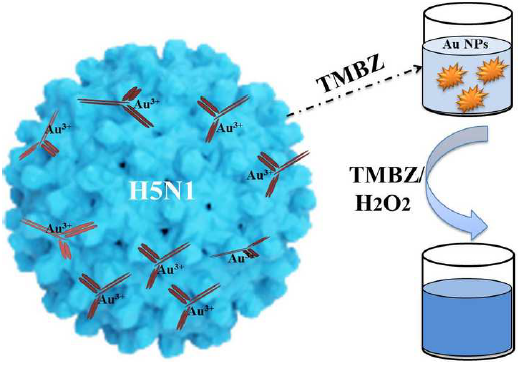

